# Human *C. difficile*-specific memory B cells encode protective IgG1 despite predominance of non-neutralizing antibodies

**DOI:** 10.64898/2025.12.09.693127

**Authors:** Sydney T. Honold, Kathleen Norris, Jordan N. May, Elizabeth J. Donald, Tyler M. Shadid, Megan Kempher, Gillian A. Lang, Jessica M. Reel, Maureen A. Cox, Jason Larabee, Jimmy D. Ballard, Kenneth Smith, Mark L. Lang

## Abstract

*Clostridioides difficile* remains a common source of nosocomial infection resulting in a wide range of clinical outcomes and for which there is no vaccine and limited therapeutic options. *C. difficile* Toxin B (TcdB)-specific IgG is the best correlate of protection against severe and recurrent disease. However, there are very few therapeutic IgG fully human monoclonal antibodies (hmAbs) that have shown efficacy thus far. We therefore hypothesized that the memory B cell (Bmem) compartment of individuals who have recovered from infection may be dominated by non-protective IgG molecules but encode some antibodies that are protective. We therefore produced hmAbs from a library of Bmem-encoded IgG1 sequences. Most TcdB-specific hmAbs displayed low affinity binding and poor neutralization of TcdB in vitro while a few hmAbs had high affinity binding and good TcdB neutralization. The results correlated with in vivo studies in which high affinity, TcdB-neutralizing hmAbs provided moderate protection of C57Bl/6 mice against *C. difficile* disease. Similar protection was observed in Tg32 mice expressing the human FcRn transgene, indicating that gut delivery of hmAb did not account for limited efficacy. Although non-protective antibody sequences dominate the repertoire, human Bmem cells may be a good source of therapeutic hmAbs for *C. difficile* treatment.

## INTRODUCTION

*Clostridioides difficile* is the most common cause of nosocomial gastrointestinal infections in the US and is responsible for 30,000 deaths annually^1^. *C. difficile* infections (CDI) also represent a significant global health issue ^2^. Colonization of the large intestine with *C. difficile* results in symptoms that range from diarrhea to life-threatening pseudomembranous colitis and toxic-megacolon. Evidence suggest that the causes of *C. difficile* associated mortality include systemic sequelae such as hepatic abscesses, ascites, acute respiratory distress, and sepsis and multi-organ failure ^3–10^.

CDI-associated pathology is attributable to secreted toxins known as TcdA and TcdB ^11,12^. These toxins enter target cells and glucosylate Rho GTPases to facilitate broad cellular damage ^13,14^ and suppression of host immunity ^15–17^. TcdA and TcdB cause systemic toxicity in several animal species ^18–21^, supporting the observations of systemic pathology in patients.

Several distinct ribotypes and strains of pathogenic *C. difficile* cause disease of varying severity ^22^. Mutation of TcdB is likely to contribute to differences in disease severity and recent studies report 8-12 distinct subtypes of TcdB ^23,24^. Infection with a hypervirulent *C. difficile* strain such as NAP1/027/BI (ribotype 027 producing TcdB2) is associated with more severe disease than a historical strain such as VPI-10463 (ribotype 003 producing TcdB1) ^25–27^. Although TcdB2 and TcdB1 share 92% sequence identity, TcdB2 is more cytotoxic than TcdB1 ^21^.

Up to 30% of individuals diagnosed with CDI will suffer from disease recurrence ^28,29^ which increases the probability of further episodes, progressively worsening pathology, and increasing mortality ^30–32^. Risk factors include antibiotic use, advanced age, suppressed immunity, and the infecting *C. difficile* strain [reviewed in ^28^]. Recurrent CDI is characterized by re-growth of bacteria that have survived antibiotic therapy or by re-infection with *C. difficile*.

*C. difficile* recurrence may also indicate that an initial CDI failed to adequately immunize the individual and confer protection against subsequent CDI. Patients with higher anti-TcdA and -TcdB serum IgG titers have lower rates of recurrence and TcdB-specific IgG is the best-known correlate of protection against *C. difficile* [reviewed in ^33^]. Bacterial load during infection correlates directly with age and inversely with toxin-neutralizing IgG titers ^34^. One mechanistic explanation for poor humoral immunity following CDI is that TcdB actively suppresses the establishment of memory B cells (Bmem) encoding protective IgG ^15,16^.

Given the potentially protective role of human IgG in CDI, human monoclonal antibody (hmAb) therapy is one option for treatment. There is currently one FDA-approved hmAb, known as Bezlotoxumab, which reduces rates of recurrence by approximately 50%, leaving room for improved therapeutics ^35,36^.

We previously isolated, sequenced and curated the memory B cell-encoded TcdB-specific from volunteers who had recovered from CDI ^17^. Initial experiments with Ig sequence-derived hmAbs suggested poor TcdB neutralization, but that study did not examine protective capacity in vivo ^17^. In the present study we have produced TcdB-specific IgG1 hmAbs from our Ig repertoire. We tested the hypothesis that although low in frequency, hmAbs with high affinity TcdB binding, in vitro neutralization, and in vivo protection against *C. difficile* are encoded by human Bmem cells. We show that from a pool of 46 hmAbs tested, only 2 displayed high affinity binding, neutralization and protection in a mouse CDI model, indicating potential for development as therapeutics.

## RESULTS

### Properties of human memory B cell-encoded TcdB-specific IgG1 monoclonal antibodies

ELISA assays were performed in which hmAbs were tested for binding to holotoxins TcdB1, TcdB2, their glucosyltransferase null mutants (TcdB1-D270N, TcdB2-D270N) referred to as D270N1 and D270N2, their respective C-terminal domains (CTD1, CTD2), and TcdB2_Δ1769-1787,_ a mutant that is enzymatically active but unable to bind host cell receptors (**Fig.1 and Fig. S1**). Assay controls included a non-specific antigen (KLH) and a pan anti-Ig capture Ab to ensure equal application of hmAbs to the plates. As shown in (**Fig. 1A**), 17-09 identified by us previously ^17^ and 42-09 displayed similar binding to all antigens, to each other, and to the therapeutic mAb Bezlotoxumab. Another hmAb, 19-09, shown as an example, exhibited weaker binding. All other hmAbs from volunteer 1009 showed similarly weak binding to the TcdB-antigens (**Fig. S1**), confirming that the memory B cell-encoded repertoire largely consists of hmAbs with low affinity binding to the toxin.

**Figure 1:**
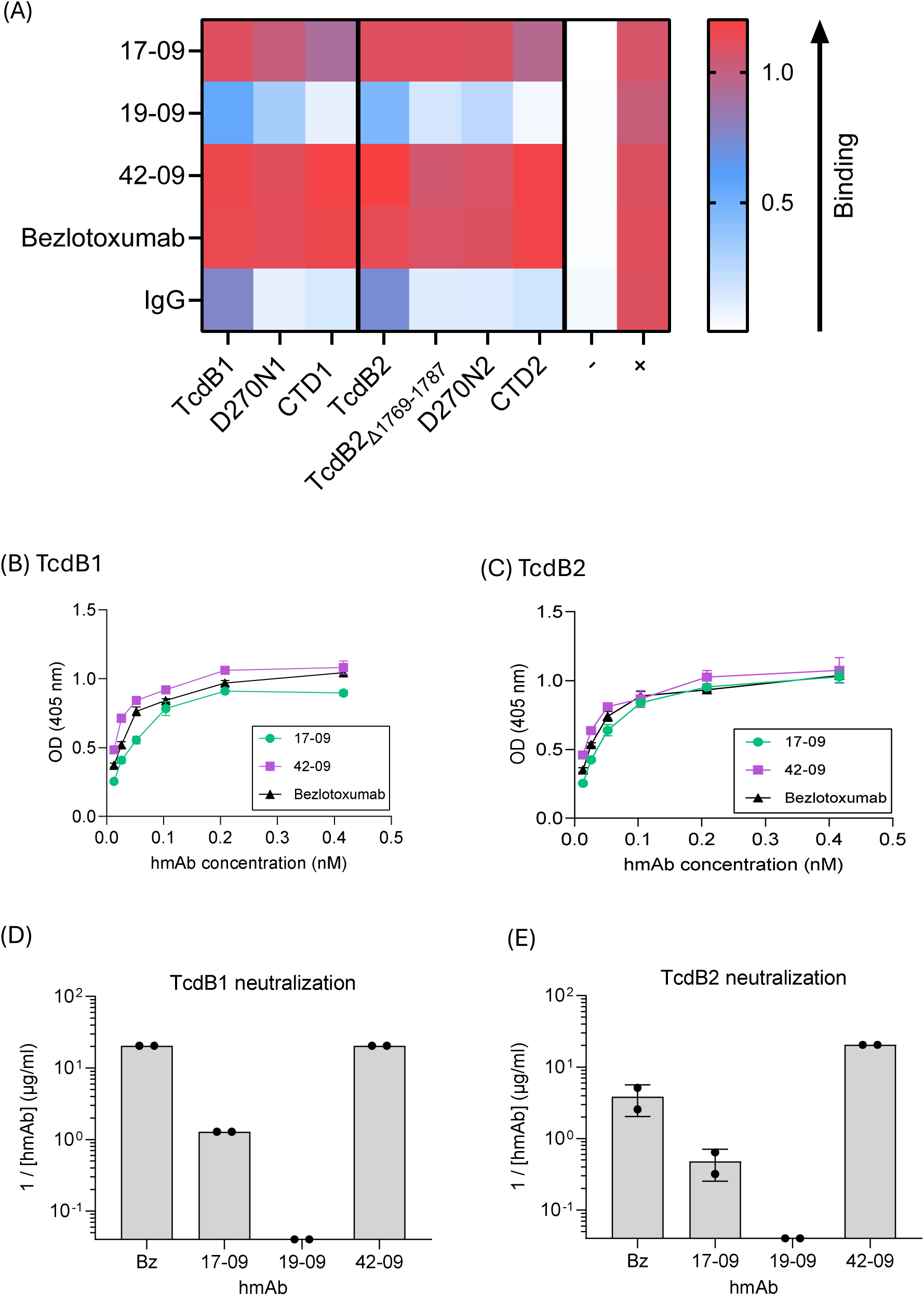
Toxin-binding properties of human Bmem-derived TcdB-specific hmAbs. ELISA analyses showing binding specificity and affinity of select hmAbs against TcdB1 and TcdB2 proteins. (**A**) Heatmap depicts binding specificity of Bmem-derived IgG1 17-09, 19-09, 42-09, Bezlotoxumab and purified human IgG against TcdB1 and TcdB2 holotoxins, the TcdB2_Δ1769-1787_ antigen, D270N mutants, and their respective CTDs. Binding to an irrelevant antigen (KLH) is also depicted. A pan anti-Ig capture ELISA was also used to demonstrate equal amounts of each hmAb in the assay. Values shown depict the mean absorbances for triplicate samples. Heatmap intensity was set automatically by GraphPad prism software. A complete table showing all IgG1 hmAbs for volunteer 1009 in the study is shown in **Fig. S1**. (**B**) Shows selected affinity binding curves for Bmem-derived 17-09, 42-09, and Bezlotoxumab against TcdB1 and TcdB2 (**C**). A series of 10 two-fold dilutions were performed in duplicate for each antigen. Graph shows mean ± SEM for duplicate samples and dissociation constants (Kd) are as indicated. Curves depicting binding to D270N and CTD proteins are shown in **Fig. S2**. (**D**) and (**E**) Depict neutralization of TcdB1 and TcdB2 respectively by the select hmAbs. Data are expressed as the reciprocal of the minimum dilution of hmAb at which 50% protection against cell rounding could be observed. Duplicate values are depicted for samples analyzed independently in different culture plates.

To further examine binding, dissociation constants (Kd) were determined by dilution of 17-09, 42-09, and Bezlotoxumab in the ELISA assays against plate-bound TcdB1 (**Fig. 1B**) and TcdB2 (**Fig. 1C**). Binding of these hmAbs and 19-09 to the D270N and CTD proteins are also shown in (**Fig. S2**). The hmAbs 17-09 and 42-09 had high affinity for TcdB-antigens with no significant differences noted in affinity for TcdB1 and TcdB2. It should be noted that Bezlotoxumab has two TcdB binding sites with different affinities ^37^ and that the assay performed herein provides only an average Kd. These experiments reveal that 17-09 and 42-09 have similar Kd values to an established therapeutic hmAb.

Similar analyses were performed using hmAbs generated from another previously infected volunteer, Patient 1008 (**Fig. S3**). Most hmAbs displayed weak binding to the various TcdB antigens used in the study with two, 7-08 and 32-08, showing strong binding to all antigens. However, both 7-08 and 32-08 failed to neutralize TcdB1 or TcdB2 in vitro (*not depicted*). These data confirm that the observations regarding hmAbs from volunteer 1009 extend to others.

Each hmAb in the study was then tested using an in vitro neutralization assay in which hmAbs were diluted against a constant TcdB1 or TcdB2 concentration applied to cultured Vero cells. Bezlotoxumab, 17-09, and 42-09 had the ability to neutralize TcdB1 (**Fig. 1D**) and TcdB2 (**Fig. 1E**). Bezlotoxumab showed superior neutralization of TcdB1 and TcdB2 as compared to 17-09, but inferior neutralization as compared to 42-09. All other hmAbs in the study failed to neutralize TcdB1 or TcdB2. These data show that the Bmem IgG1 compartment largely encodes non-neutralizing IgG1, (44/46 hmAbs tested), but a minority (2/46 tested) show neutralization comparable to the well-established therapeutic hmAb, Bezlotoxumab.

### Memory B cell-encoded hmAbs partially protect C57Bl/6 mice from a TcdB2-secreting *C. difficile* strain

The ability of 17-09, 19-09, 42-09, and the therapeutic hmAb Bezlotoxumab to protect C57Bl/6 mice against CDI was tested as outlined (**Fig. 2A).** Mice in the treatment groups were administered hmAb 24 hours prior to oral gavage with R20291 *C. difficile* spores. Infected, but untreated mice demonstrated statistically significant weight loss as compared to uninfected controls (**Fig. 2B**). Significant weight loss was observed in all experimental groups, although timing of peak weight loss and recovery varied slightly between experiments. Bezlotoxumab and hmAb 17-09 minimized weight loss, whereas the non-neutralizing hmAb19-09 did not. Comparison of weights at days 0 and 2 for each infected mouse confirmed these observations (**Fig. 2C)**. An ANOVA analysis also confirmed that on days 3 and 4, weight loss was significantly reduced by Bezlotoxumab and 17-09 but not 19-09 (as compared to infected controls) (**Fig. 2D, left panel**). The hmAb 42-09 was produced later in the study and tested in a separate experiment **(Fig. 2D, E).** Treatment with hmAb 42-09 significantly reduced weight loss (at days 2 and 3) as compared to infected but untreated mice (**Fig. 2D, 2E**). These data therefore demonstrate that 17-09 and 42-09 show efficacy in protection of C57Bl/6 mice against a TcdB2-secreting *C. difficile* strain.

**Figure 2:**
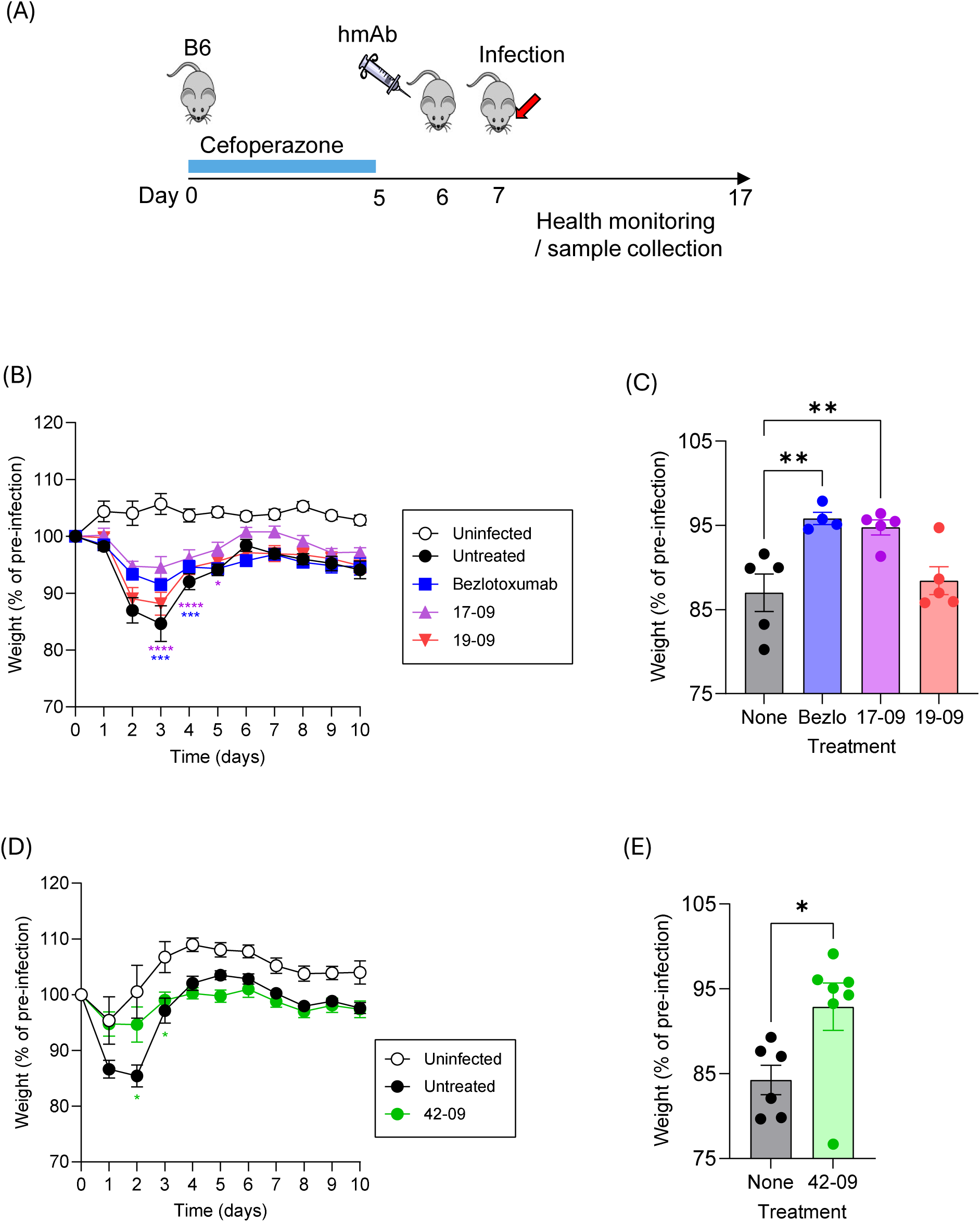
Bmem-encoded hmAbs 17-09 and 42-09 partially protect B6 mice from *C. difficile* disease. (**A**) B6 mice were given Cefoperazone in drinking water as indicated before withdrawal and treatment with hmAbs as indicated on day 6. Live R20291 *C. difficile* spores were administered by oral gavage on day 7 before daily health monitoring. (**B**) Shows mean ±SEM changes in weight across 10 days in each experimental group (n=5 / group). Tested hmAbs include Bezlotoxumab, 17-09, and 19-09. A two-way ANOVA with Dunnett’s multiple comparison post-test was performed (***, p<0.001, **** p<0.0001). (**C**) Shows mean ±S.D. relative weights of each group at day 2. A one-way ANOVA with Dunn’s post test was performed (**, p<0.01). (**D**) Mean ±SEM changes in weight across 10 days in each experimental group (n=5 / group) for parallel experiment testing 42-09. (**D**) Shows mean ±S.D. relative weights of each group on day 2. A two-tailed t-test was performed comparing weight loss at day 2 (**, p<0.01). Data are representative of two similar experiments.

In a further experiment, Bezlotoxumab and the hmAbs 17-09, 42-09 and 19-09 were administered 24 hours prior to oral gavage with R20291 *C. difficile* spores. Colonic tissue was harvested 3 days after infection and prepared for analysis (**Fig. 3**). Composite disease scores equally weighted immune cell infiltration, epithelial cell damage, edema, and weight loss. The results showed that infection but not Cefoperazone treatment significantly increased the disease score and that the small non-significant increase caused by the antibiotic was due to edema (**Fig. 3A, B**). Although they limited weight loss, none of the hmAbs tested showed a significant reduction in overall disease score, including Bezlotoxumab and hmAbs 17-09 and 42-09 (**Fig. 3A, B**). This indicates that hmAb treatment did not significantly improve gut pathology during infection.

**Figure 3:**
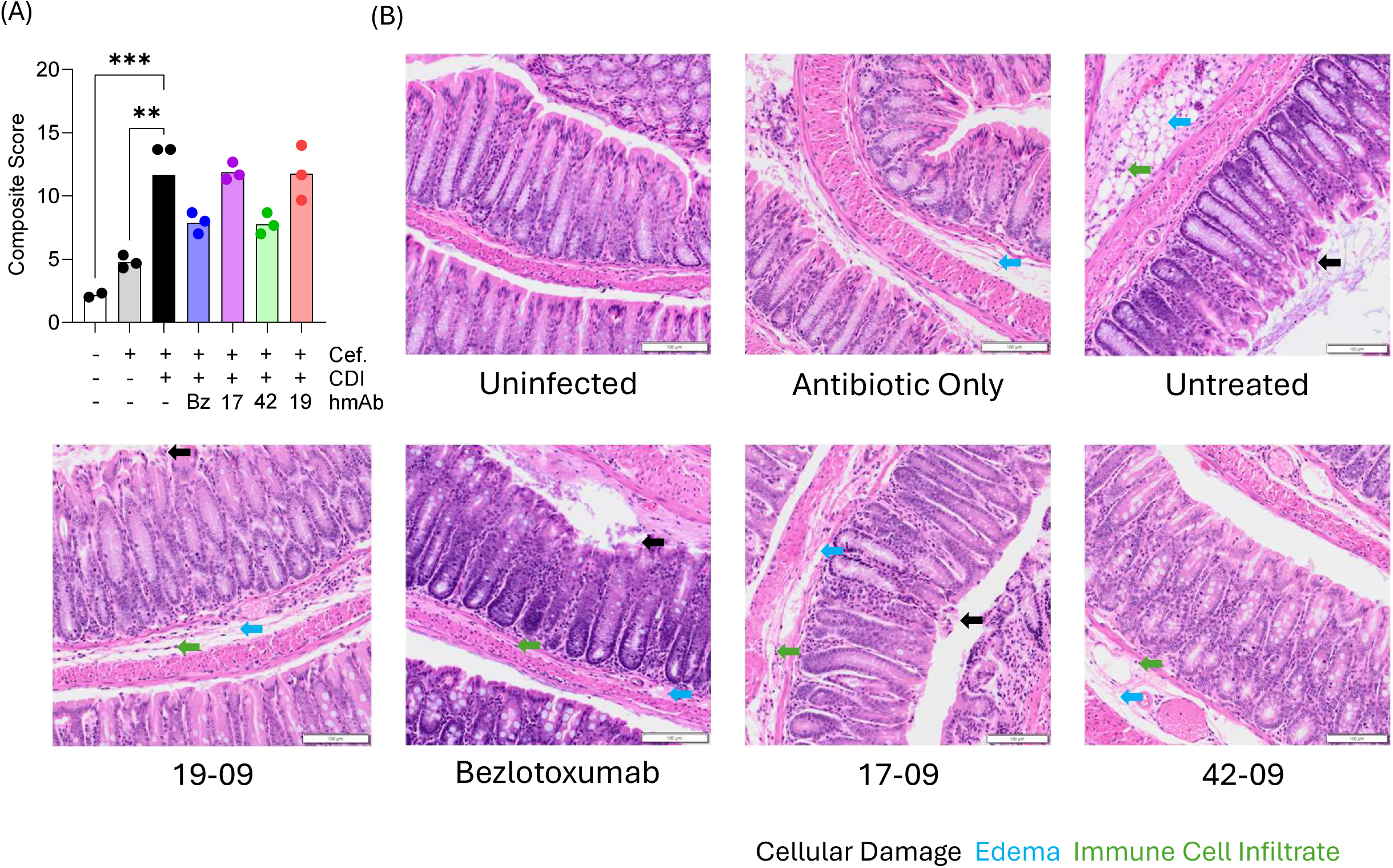
Modest impact of Bmem-encoded hmAbs 17-09 and 42-09 on colonic pathology. H&E analysis of full length ‘Swiss-rolled’ colons was performed 3 days post CDI as described in methods. (**A**) Shows composite scores for disease severity (n=3 / group). Data were analyzed by One-way ANOVA with Holm-Sidak’s multiple comparison test (**, P<0.01, ***, P<0.001). (B) Representative images of H&E-stained colon tissue for each treatment group are shown.

### The efficacy of hmAb delivery to the gut lumen does not explain the limited protection in vivo

To ensure that moderate protection of B6 mice with 42-09 was due to limited efficacy rather than delivery to the gut lumen, we tested its ability to protect Tg32 mice against CDI (**Fig. 4A).** The FcRn receptor facilitates transport of IgG to the gut lumen and increases the half-life of IgG ^38^. Tg32 mice (FcRn^−/-^) express the human FcRn transgene and are used for testing hmAbs in vivo (reviewed in ^39^). Infected, untreated mice demonstrated statistically significant weight loss as compared to uninfected controls (**Fig. 4B**). Treatment with hmAb 42-09 showed no significant effect on weight loss over the experiment as assessed by two-way ANOVA (**Fig. 4B**) but by matched pairs t-test limited weight loss on day 3 (**Fig. 4C**). Male and female mice were included in this study and disease course in controls or hmAb-treated groups did not differ. Bacterial burden as expected was not altered by 42-09 treatment (**Fig. 4D**). To confirm delivery of hmAb to the gut in Tg32 mice, ELISA analysis of fecal pellets was performed. Bezlotoxumab was the most abundant hmAb available for experiments and was used to show delivery to the gut lumen consistent with FcRn protecting IgG from degradation and / or boosting delivery of hmAbs to the gut (**Fig. 4E**). These data therefore confirm that hmAb 42-09 offers modest protection of Tg32 and B6 mice against a TcdB2-secreting *C. difficile* strain. Furthermore, limitations on delivery to the gut or hmAb half-life do not explain their modest efficacy *in vivo*.

**Figure 4.**
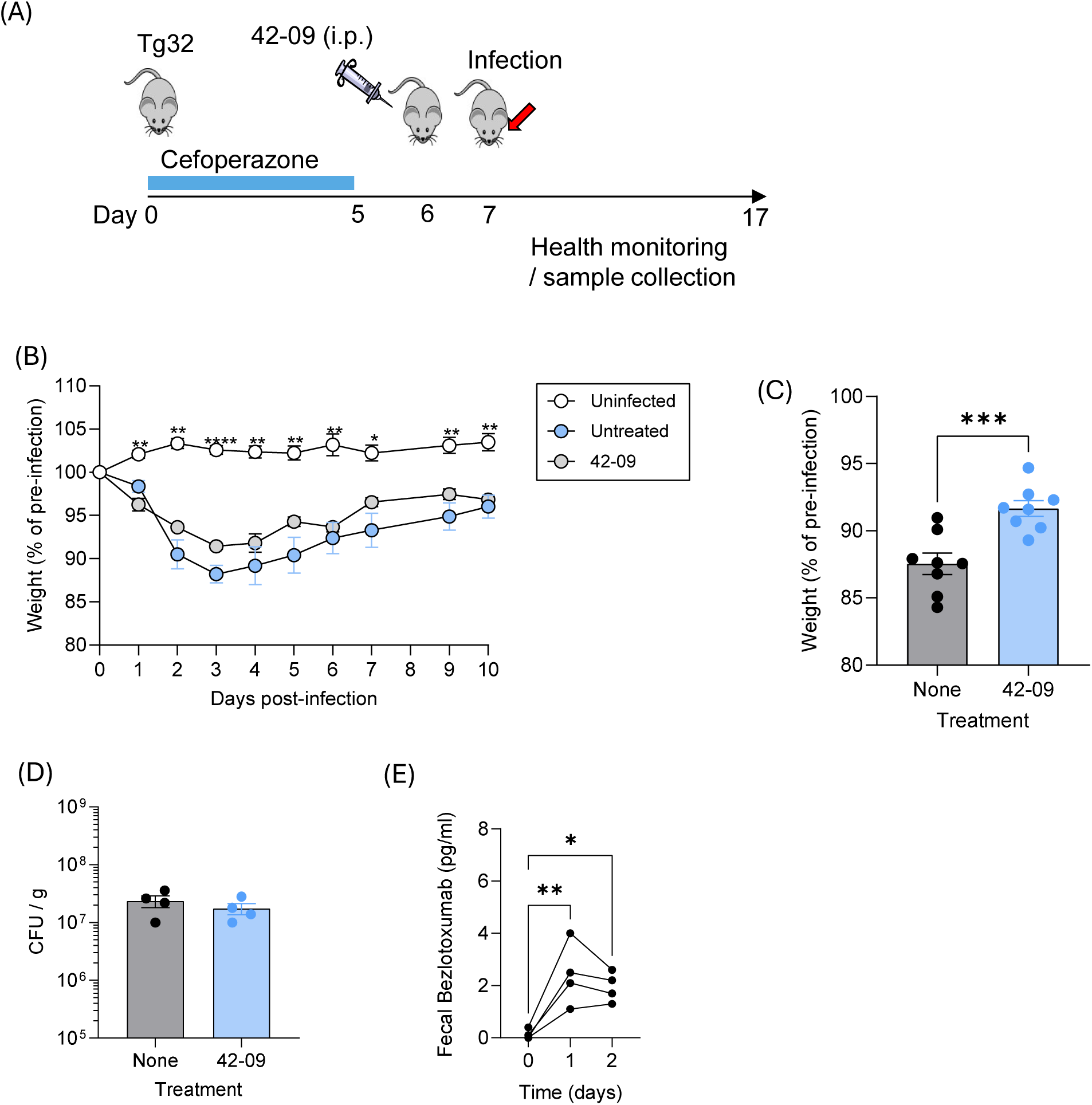
Bmem-encoded hmAb 42-09 partially protects Tg32 mice from *C. difficile* disease. **(A)** Male and female Tg32 mice were given cefoperazone in drinking water before withdrawal and treatment with 42-09 as indicated on day 6. Live R20291 *C. difficile* spores were administered by oral gavage on day 7 before daily health monitoring. (**B**) Shows mean ±SD changes in weight in each experimental group (n=11 for uninfected, n=5 for untreated and 4 for hmAb 42-09-treated) and is representative of 2 similar experiments. Significance was determined using two-way ANOVA and Dunnett’s multiple comparison test. **, p<0.01, ****, p<0.0001. (**C**) Shows mean ±S.D. minimum weights achieved (p<0.001 by matched pairs t-test). Pooled data from two independent experiments is shown for. (**D**) Shows fecal CFU counts. (**E**) Tg32 mice were injected by the i.p. route with 100 µg Bezlotoxumab before its detection and measurement in fecal pellets by ELISA. (*, p<0.05, **, p<0.01 by one way ANOVA and Tukey’s post-test).

## DISCUSSION

In this study we have examined the functional TcdB-specific human Bmem repertoire following *C. difficile* infection. Of the 46 IgG1 sequences produced as hmAbs, only four showed strong binding affinity for TcdB and related antigens and only two neutralized TcdB1 and TcdB2 in vitro. The hmAbs that neutralized TcdB1 and TcdB2 in vitro partially protected B6 and Tg32 mice in a CDI infection model. The hmAbs 17-09 and 42-09 performed similarly to Bezlotoxumab, limiting CDI-induced weight loss caused by a TcdB2-secreting *C. difficile* strain but had limited impact on pathology in the colon.

This study therefore shows that the human CDI-specific antibody response following infection is mostly non-protective against disease. Consistent with this, achieving TcdB2 neutralization by vaccination has proven difficult. And has required engineering of the TcdB2 protein to expose protective B cell epitopes ^40^. However, hmAbs with some therapeutic potential may be present in the Bmem-encoded antibody repertoire following infection. Screening the Bmem compartment of individuals following CDI could lead to identification of a wider array of therapeutic hmAbs.

It is unknown if individuals with recurrent disease have fewer TcdB-specific Bmem cells, or a lower frequency of Bmem cells that encode neutralizing IgG than individuals who only suffered a single infection. A larger-scale study would be necessary to fully answer this question. The function of the Bmem cell and their capacity to differentiate into functional plasma cells that can secrete IgG (or neutralizing IgG) is also unknown. In our previous studies, we did observe that polyclonal stimulation of Bmem cells resulted in differentiation of new TcdB-specific plasmablasts that could be detected by ELISPOT^41^. Furthermore, serum from the individuals in this study can neutralize TcdB, showing that individuals can accumulate protective plasma cell-derived TcdB-specific IgG ^17^. However, those observations do not reveal whether those plasma cells developed from the Bmem pool. There are therefore numerous avenues to explore and understand the humoral immune response of the previously infected patient.

Bezlotoxumab is reported to reduce disease recurrence in humans by around 50%^35 36^. The hmAbs 17-09 and 42-09 could potentially limit disease recurrence and future studies will address this question. One hypothesis for why these hmAbs could limit disease recurrence is that by neutralizing TcdB they could remove TcdB-induced suppression and allow a better host Bmem response to the pathogen. In a previous study, we demonstrated that TcdB2 impacts germinal center reactions and Ig class switch by altering CXCR4-dependent B cell migration^16^. The hmAbs in this study could be tested for their ability to block TcdB subversion of host humoral immunity. However, it may also be hypothesized that highly efficient TcdB neutralization would remove the source of antigen needed to immunize the host and mount an effective anti-TcdB response. While testing these hypotheses experimentally would be challenging, the observation that hmAbs 17-09 and 42-09 were not fully protective does suggest the possibility of incomplete TcdB2 neutralization in the mouse model of CDI. This could perhaps be exploited to achieve a good balance between intoxication and antigen availability. Arguably, it could then be determined whether protective hmAbs can limit immunosuppression during a primary infection and allow host immunity to mitigate a recurrent infection.

Our study also provokes questions regarding how hmAbs 17-09 and 42-09 protect since impact on colon pathology was limited. This is consistent with recent studies testing a TcdB-specific hmAb ^42^. The results suggest that hmAbs potentially protected against systemic effects of TcdB2 or against sepsis caused by gut commensals. Determination of the TcdB2 epitopes and hmAb binding sites may be informative. Also, understanding the relatedness of new hmAbs as they are discovered may be important. For example, if there is some degree of convergent evolution of protective hmAbs in that they exhibit the same gene usage or similar critical sequences in the variable regions, then new hmAbs could be synthesized without needing to access patient samples. Such analyses may await the discovery of several new neutralizing hmAbs before a pattern emerges.

Discovery of new TcdB2-neutralizing hmAbs also prompts the question of whether therapies should be developed that consist of hmAb cocktails to provide greater coverage against toxinotypes. There are at least 8 and perhaps 12 distinct toxinotypes, and greater coverage than that provided by current therapeutic mAbs may be needed. Expression of hmAbs as IgG2 rather than IgG1 may be worth testing, as may engineering of constant regions to enhance FcRn-mediated delivery of IgG to the gut.

In summary this study shows the utility of screening the human Bmem repertoire to discover therapeutic hmAbs to treat CDI and to assess the protection potential in the previously-infected individual.

### Limitations of study

The in vivo experiments focused on a single *C. difficile* strain and a small number of test hmAbs. To fully understand the utility of Bmem-derived hmAbs, larger scale studies may be required. It should also be noted that the disease course is shorter in mice than in humans and probing the impact of hmAbs on toxin-suppressed humoral immunity and disease recurrence may be challenging.

## MATERIALS AND METHODS

### Sex as a biological variable and experimental model

Female C57Bl/6 mice at 6-8 weeks of age were purchased from Charles River Laboratories (Bethesda, MD, Cat # model 027) or Jackson Laboratories (Stock #: 000664). Mice lacking the endogenous FcRn and expressing the human FcRn transgene (Tg32) were purchased from Jax.org (Stock #: 014565). Tg32 mice were bred in house and genotyping confirmed by PCR (performed by Transnetyx, Cordova, TN). Both male and female Tg32 mice were used for experiments. We have never observed sex related differences in the *C. difficile* infection model.

### Purification of TcdB and TcdB-derived proteins

TcdB1, TcdB2, TcdB2_Δ1769-1787_, D270N1, D270N2, were expressed in *Bacillus megaterium* (MoBiTec, Göttingen, Germany) and purified by nickel affinity chromatography (GE Life Sciences, Boston, MA) as previously described ^40^. CTD1, and CTD2 were expressed in *Escherichia coli* BL21star DE3 (Invitrogen, Carlsbad, CA). TcdB1 and TcdB2 refer to bioactive holotoxins. The D270N point mutation renders the toxin glucosyl-transferase null ^43,44^. TcdB2_Δ1769-1787_ has a 19 amino acid deletion in the receptor binding domain that negates cell binding ^40^. CTD1 and CTD2 refer to C-terminal domain proteins (AA165-2366) encapsulating the C-terminal CROP regions of the toxin (AA1834-2366) ^45^. Purity and integrity were confirmed by SDS-PAGE and each batch of TcdB1 and TcdB2 was tested for toxicity using a CHO cell killing assay ^46^.

### Expression and Purification of full-length hmAbs

TcdB-specific hmAb sequences were acquired as described in ^17^ and produced as full-length hmAbs as described previously ^47^. Selected hmAbs that showed high binding and neutralization capacity were synthesized in bulk using the TurbCHO Antibody Expression service offered by GenScript (Piscataway, NJ). Commercial grade Bezlotoxumab was purchased from MedChemExpress (CAT: HY-P9929, Lot: 251891) (Monmouth Junction, NJ) and produced by Genscript. The Human IgG Isotype Control (CAT:02-7102, Lot: YI384682) was from Invitrogen (Carlsbad, CA).

### *C. difficile* spore preparation

*C. difficile* R20291 spores were prepared and isolated as previously described ^48,49^. Briefly, a single colony grown on Brain Heart Infusion (BHI) + Taurocholic acid (TCA) was used to inoculate 2 ml of Columbia Broth and grown overnight at 37⁰C anaerobically. The next day, the 2 ml culture was used to inoculate 40 ml of Clospore^TM^ media and cultured anaerobically at 37⁰C for 5-7 days. Spores were collected by centrifugation (4,000 rcf, 4⁰C, for 20 minutes) and washed with cold water at least three times and until the supernatant was clear. Finally, the spore pellet was resuspended in 1 ml sterile water and placed in a glass vial and stored at 4⁰C. Before infection, *C. difficile* spores were enumerated by plating on TCCFA (Taurocholate Cycloserine Cefoxitin Fructose Agar). A pre-calculated concentration of spore inoculum was heated at 65°C for 20 min, then allowed to cool for 5 min at room temperature before use.

### *C. difficile* infection and hmAb treatment

Mice were housed in sterile cages with sterile food. Mice were provided with Cefoperazone sodium salt (Millipore Sigma, St. Louis, MO) in distilled drinking water (0.5 g / l) for 5 days, followed by 2 days of distilled water (Thermofisher, Waltham, MA). Mice were infected by oral gavage with 1 × 10^5^ heat-treated *C. difficile* R20291 spores. *C. difficile*-associated pathology was assessed by monitoring daily weights, and other clinical signs such as lethargy, hunched posture, and diarrhea. Animals were euthanized if the weight loss reached 20.0% or the mice were moribund ^50^. Bacteria were quantified on day 3 post-gavage by homogenizing fecal pellets with 1X PBS, serially diluted, plated on TCCFA and cultured under anaerobic conditions at 37°C. CFUs were counted within 24 hours ^50^. Where indicated mice were injected by the intraperitoneal (i.p.) route with 100 µg of hmAb in sterile PBS 24 hours before *C. difficile* challenge.

### ELISA assays

#### (1) hmAb reactivity to TcdB-type antigens

Nunc MaxiSorp Enzyme-linked Immunosorbent Assay (ELISA) 96-well plates (Thermo Scientific, Waltham, MA) were coated overnight at 4°C with carbonate coating buffer (pH 9.2) and the indicated protein (TcdB1, TcdB2, D270N1, D270N2, TcdB2_Δ1769-1787_, CTD1, or CTD2) at a final concentration of 10 µg/mL. Wells were washed 4 times with PBS-T (1x PBS, 0.05% Tween) using BioTek 405 Microplate washer (Winooski, VT) and blocked with 200 µl of 1% bovine serum albumin (BSA) in PBS-T (blocking buffer) for 2 hours at room temperature. The plates were then washed 4 times as before with PBS-T. Wells were incubated with specified hmAb diluted in blocking buffer to a final concentration of 10 µg/mL for 2 hours at room temperature. Plates were then washed 4 times as before with PBS-T. Wells were incubated with HRP-anti human IgG (Jackson ImmunoResearch, 1:2500 in blocking buffer) for 1 hour at room temperature in light-protected conditions. To develop, plates were washed 4 times as before with PBS-T, and 90 µl of ABTS (pre-warmed to room temperature) was added to each well for 5 minutes under light-protected conditions. Reaction was stopped by additions of 110 µl of stop solution (10% SDS w / v ddH_2_O) to each well. Samples absorbance ar at 405 nm was measured using a Molecular Devices SpectraMax ABS plate reader (San Jose, CA). Where indicated selected hmAbs were subjected to a series of ten 2-fold dilutions starting at 10 µg/mL (19-09) or 1 µg/mL (17-09, 42-09, and Bezlotoxumab) final concentration in the assay. The absorbance values obtained for each hmAb concentration were applied to a curve-fitting analysis to calculate Ab affinities (K_D_).

#### (2) Detection of hmAbs in mouse fecal samples

Fecal pellets were suspended in proteinase inhibitor solution to 10% w/v (EMD Millipore Corp., Burlington, MA) and incubated overnight at 4°C. The pellets were then homogenized by shaking for 5 seconds followed by vortexing until homogenized (typically 15-30 seconds). Samples were then centrifuged at 15,000 *rcf* for 30 minutes at 4°C before collecting supernatants. Samples were stored at -20°C until required. For ELISAs, a 4-fold dilution of the sample was applied to pre-coated and blocked ELISA plates. Detection of human IgG was as described.

### Cell culture and *in vitro* neutralization

Vero cells were cultured in 96 well plates until 50% confluence was achieved. TcdB1 or TcdB2 were applied to the cultures at final concentrations of 0.6 and 2 ng/ml respectively. These concentrations have been determined to cause 95% cell rounding in the assay. TcdB was applied alone or after pre-mixing with hmAbs for 30 min at 37°C. After 24 hours, microscopy was used to determine the concentration of hmAb at which 50% of Vero cells had adopted a rounded morphology. Duplicate culture plates were used for every experiment and blinded analyses were performed.

### Histology

Mice were euthanized on day 3 post-oral gavage. Each colon was excised, cut lengthwise, and opened to lay flat. A cotton swab dipped in 1x PBS was used to gently remove any feces and to help position the tissue, which was then gently rolled starting from the anal junction end. Once rolled, the tissues were pierced with a 30-gauge needle, placed in tissue cassettes, and fixed in Modified Davidson’s fixative (64% DI water, 20% ethanol, 6% glacial acetic acid, 10% formaldehyde). Paraffin sections (5 µm thick) were mounted on slides and stained with hematoxylin and eosin (H&E) (performed by Excalibur Pathology Inc., Norman, OK). Slides were imaged on an Olympus SlideView VS200 (Center Valley, PA) at 20X using the Default Brightfield Scan protocol. Images were exported as .tif files and uploaded to HistoWiz.com (Long Island City, NY) for blinded disease index measurement by a board-certified pathologist. Overall disease index of the colonic tissue was quantified by scoring immune cell infiltration, epithelial damage, and edema on a scale of 0 – 4 using a previously published rubric ^51^. A composite disease score was calculated for each mouse in the study that included tissue pathology and weight loss. Immune cell infiltration, epithelial damage, edema, and weight loss were weighted equally.

### Experimental Design

Experiments were designed with prior consideration of statistical power and were repeated to demonstrate reproducibility. Figure legends provide information on group size (n), statistical test used, number of experiments performed, and whether data shown are representative or pooled. Blinded analyses were performed for toxin neutralization assays and histology images.

### Statistics

Data were analyzed using GraphPad Prism (Version 9.1.1, La Jolla, CA). A two-tailed t-test or a Mann-Whitney U test, and one-way ANOVA with Dunnett’s or Holm-Sidak’s multiple comparison tests were used for statistical analysis between two and multiple experimental groups respectively. Where indicated, paired t-tests were used. A Two-way Repeated Measure ANOVA with Tukey’s multiple comparisons test was used to determine statistical significance in weight loss measured at multiple time points. A p value of 0.05 or less was considered statistically significant.

### Study Approval

This study was performed in accordance with the recommendations of the Guide for the Care and Use of Laboratory Animals of the National Institutes of Health. All animal procedures were therefore approved by the Institutional Animal Care and Use Committee at the University of Oklahoma Health Sciences Center (OUHSC) (protocol 23-037-ACHIU). Inhalational anesthesia using a 4% isoflurane / 96% medical air mixture dispersed through a precision vaporizer was used for ear tagging and tissue collection for genotyping. Anesthesia was not used for oral gavage, weight measurements, fecal pellet collection, or intraperitoneal injections.

### Dataset availabilit

Sequences were submitted to the NCBI Sequence Read Archive (SRA) and the project is registered with the BioProject database under SRA accession: PRJNA601978 (http://www.ncbi.nlm.nih.gov/bioproject/601978). The accession numbers for the data are: SAMN13878387 (1008); SAMN13878388 (1009); SAMN13878389 (1013).

## ACKNOWLEDGEMENTS

The research reported herein was funded by NIH grant U19 AI174994 to MLL, JDB and MAC, and RO1 AI185203 to JL. Generation of antibody gene sequences prior to the experiments reported herein was supported by a Presbyterian Health Foundation of Oklahoma City Team Science Award and the Oklahoma Shared Clinical and Translational Resources (OSCTR) (U54GM104938) and reported in 10.1172/jci.insight.138137 (reference 17 in this article). The content is solely the responsibility of the authors and does not necessarily represent the official views of the National Institutes of Health.

## CONTRIBUTIONS

STH: Experimental Design; Project coordination (hmAb production); Experimental (binding assays, histology); Data analysis; Writing (MS text, editing, figure preparation).

KN: Experimental (hmAb production and quality control).

JNM: Experimental (challenge assays, fecal ELISA); Data analysis (challenge assays, fecal ELISA).

GAL: Experimental (training, assay design, Tg32 colony); Writing (editing).

JMR: Experimental (colon tissue preparation).

TMS: Experimental (spore prep, oral gavage, CFU counts, protein expression and purification).

MK: Experimental (protein engineering, protein expression and purification).

ED: Experimental (neutralization assays); Data analysis (neutralization assays).

JL: Assay design and supervision (neutralization assays); Resources (neutralization assays).

MAC: Resources and supervision (tissue preparation).

JDB: Conceptualization and supervision (antigen design, expression, purification, challenge assays); Resources (antigen design, expression, purification, challenge assays).

KS: Conceptualization / project design /resources (hmAb production).

MLL: Project conceptualization and design; supervision; resources; data analysis; writing; editing; figure preparation.

